# Mining folded proteomes in the era of accurate structure prediction

**DOI:** 10.1101/2021.08.24.457439

**Authors:** Charles Bayly-Jones, James Whisstock

**Affiliations:** Department of Biochemistry and Molecular Biology, Monash University, Clayton, Victoria, Australia; Biomedicine Discovery Institute, Faculty of Medicine, Nursing and Health Sciences, Monash University, Clayton, Victoria, Australia

## Abstract

Protein structure fundamentally underpins the function and processes of numerous biological systems. Fold recognition algorithms offer a sensitive and robust tool to detect structural, and thereby functional, similarities between distantly related homologs. In the era of accurate structure prediction owing to advances in machine learning techniques, previously curated sequence databases have become a rich source of biological information. Here, we use bioinformatic fold recognition algorithms to scan the entire AlphaFold structure database to identify novel protein family members, infer function and group predicted protein structures. As an example of the utility of this approach, we identify novel, previously unknown members of various pore-forming protein families, including MACPFs, GSDMs and aerolysin-like proteins. Further, we explore the use of structure-based mining for functional inference.

## Main

Knowledge of a proteins’ structure is a powerful means for the prediction of biological function and molecular mechanism^1,2^. Accordingly, powerful pairwise fold recognition tools such as DALI^3^ have been developed that permit searching of known fold space in order to identify homology between distantly related structurally characterised proteins. These approaches can identify homologous proteins even when primary amino acid sequence similarity is not readily detectable. This method is particularly useful when a protein of no known function can be flagged as belonging to a well characterised fold class (e.g. Rosado et al.^4^).

A key and obvious limitation of using fold recognition to infer function is that the structure of the protein of interest needs to first be determined. Now, however, in the era of accurate protein structure prediction^5,6^, it is possible to build a reasonably accurate library comprising representative structures of all proteins in a proteome^7–9^ (**Fig 1a-c**). One utility of such a resource, is that fold recognition approaches for prediction of function can now be applied to any protein (**Fig 1d, e**).

**Figure 1.**
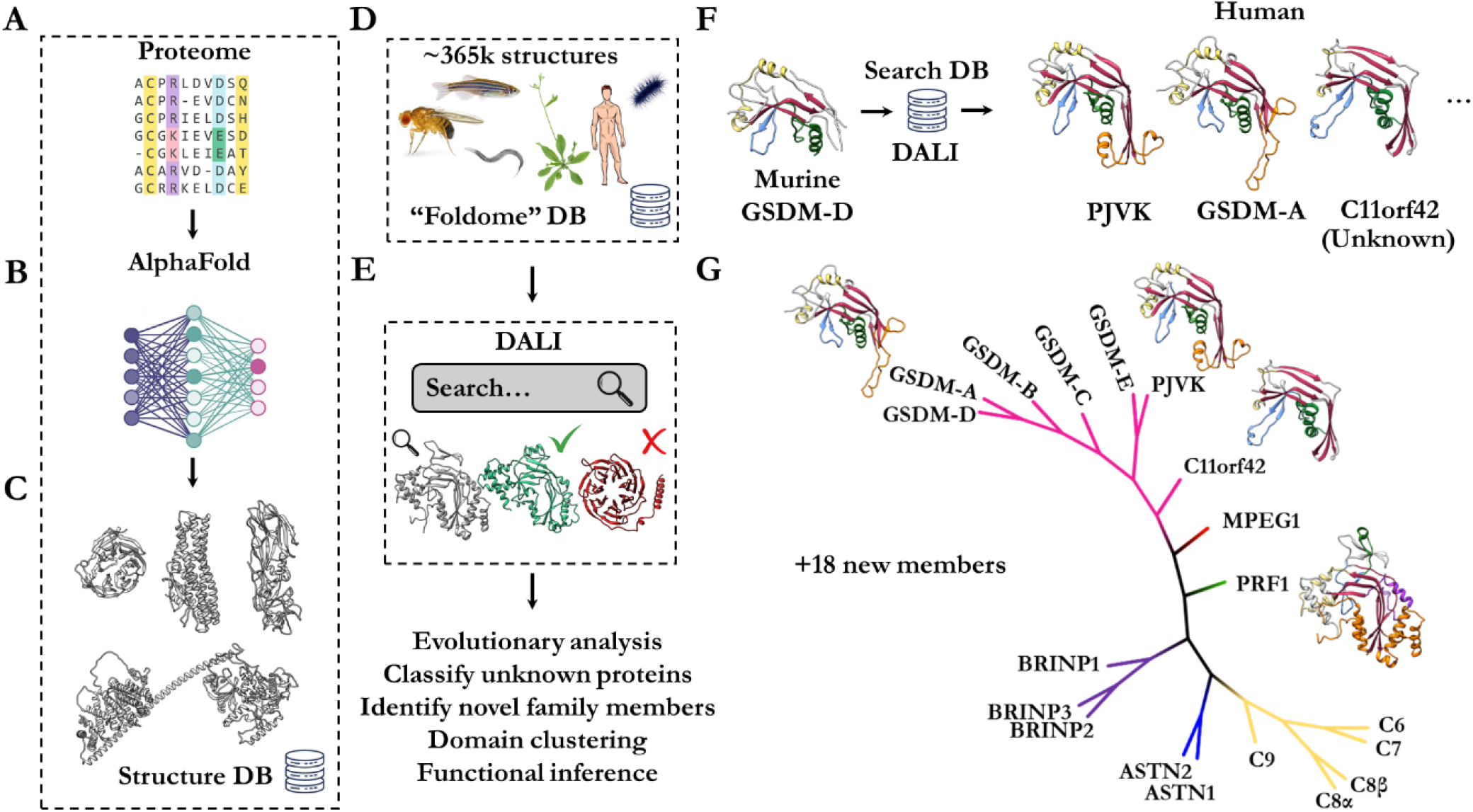
Conceptual overview of structure-guided fold recognition against the AlphaFold database. **a**. Entire proteome sequence databases are converted by (**b**) machine-learning methods into high-accuracy (**c**) structural model predictions. **d**. Curation of these structural databases into a unified resource of structures – the “foldome”. **e**. DALI based fold matching to perform functional inference, identification of unknown members and structural classification. **f**. Searching the foldome in (**d**) for matches of murine GSDM-D N-terminal domain yields MACPF/GSMD family members, including C11orf42 – an unknown member of the GSDM family. **g**. Example phylogenetic analysis of perforin-like proteins identified from the foldome by DALI. Alignment of the central MACPF/GSDM fold is possible by extracting domain boundaries based on AlphaFold prediction allowing specific comparison between members without interfering ancillary domains.

To investigate the utility of this approach we used established bioinformatic tools to mine the foldome. We constructed a locally hosted DALI database of all protein structures predicted by AlphaFold, covering humans to flies to yeast (**Fig 1d, e**). We then began mining the whole database using a probe structure representing a well characterised protein superfamily (in this case the perforin-like superfamily of pore forming immune effectors). This approach readily yields functional insight into previously uncharacterised proteins (**Fig 1f, g**; **Supp Table 1**).

For example, structure-based mining identified all known perforin / GSDM family members, but also identified a likely new member of the GSDM pore-forming family in humans, namely C11orf42 (uniprot Q8N5U0 - a protein of no known function). Remarkably there is only 1% sequence identity between the GSDMs and C11orf42 despite predicted conservation of tertiary structure. In humans, C11orf42 is expressed in testis and is highly expressed in thyroid tumours^10^. Moreover, CRISPR screens^11–13^ identified C11orf42 as contributing to fitness and proliferation in lymphoma, glioblastoma and leukaemia cell lines (BioGRID gene ID 160298)^14^.

Identification of C11orf42 as a likely GSDM family member permits several useful predictions. Owing to the presence of a GSDM fold, we postulate that C11orf42 may share GSDM-like functions such as oligomerisation and membrane interaction. Unlike other GSDMs^15,16^, however, inspection of the predicted structure suggests that C11orf42 lacks membrane penetrating regions entirely. These data imply that C11orf42 may have lost the ability to perforate lipid bilayers and instead may function as a scaffold of sorts, as has been postulated for members of the perforin superfamily^17^.

We next expanded our analysis to all proteomes covering 356,000 predicted structures; these computations take ∼24 hours on a 16 CPU Intel i7 workstation. We identified roughly 16 novel perforin-like proteins across the twenty-one model organisms covered by the AlphaFold database (**Fig 1g**; **Supp Table 1**; **Supp Data 1**). Domain boundaries defined by the structure prediction were identified manually and the perforin/GSDM-like domains were aligned based on fold. We constructed a phylogenetic tree based on the structure-constrained multiple sequence alignment (MSA), suggesting that C11orf42 is potentially related to the precursor of the common GSDMs. Curation of sequences based on predicted structures, such as this, may enable further, more comprehensive evolutionary analyses.

We next decided to perform these foldome-wide searches for several other pore-forming protein families, identifying new members of aerolysins, lysenins, cry1 toxins and more (**Supp Fig 1**; **Supp Table 1**; **Supp Data 1**). Members of these toxin families have applications in next-generation sequencing (both DNA/RNA^18,19^ and polypeptide^20–23^), as well as agricultural applications in crop protection. We anticipate the new members of these families to be of utility in translational research programs. Remarkably, many of the hits we identified suggest the unexpected presence of pore-forming protein families that were previously thought to be entirely absent in the selected phyla – for example aerolysin-like proteins in *Drosophila, C. elegans*, yeast and zebrafish.

To further assess the utility of the database in functional inference, we curated a subset of the human proteome corresponding to uncharacterised proteins of unknown function (**Fig 2 a-c**). These proteins are largely unannotated, lacking both domain and functional descriptions. We pruned all regions of the predicted structures to have pLDDT (per-residue confidence score) greater than 70 and discarded models for which fewer than 100 residues remained. These became the probe structures for iterative searches against the whole human foldome to identify known proteins with assigned domains and function. We provide these as supplementary data for the convenience of the reader (**Supp Data 2**).

**Figure 2.**
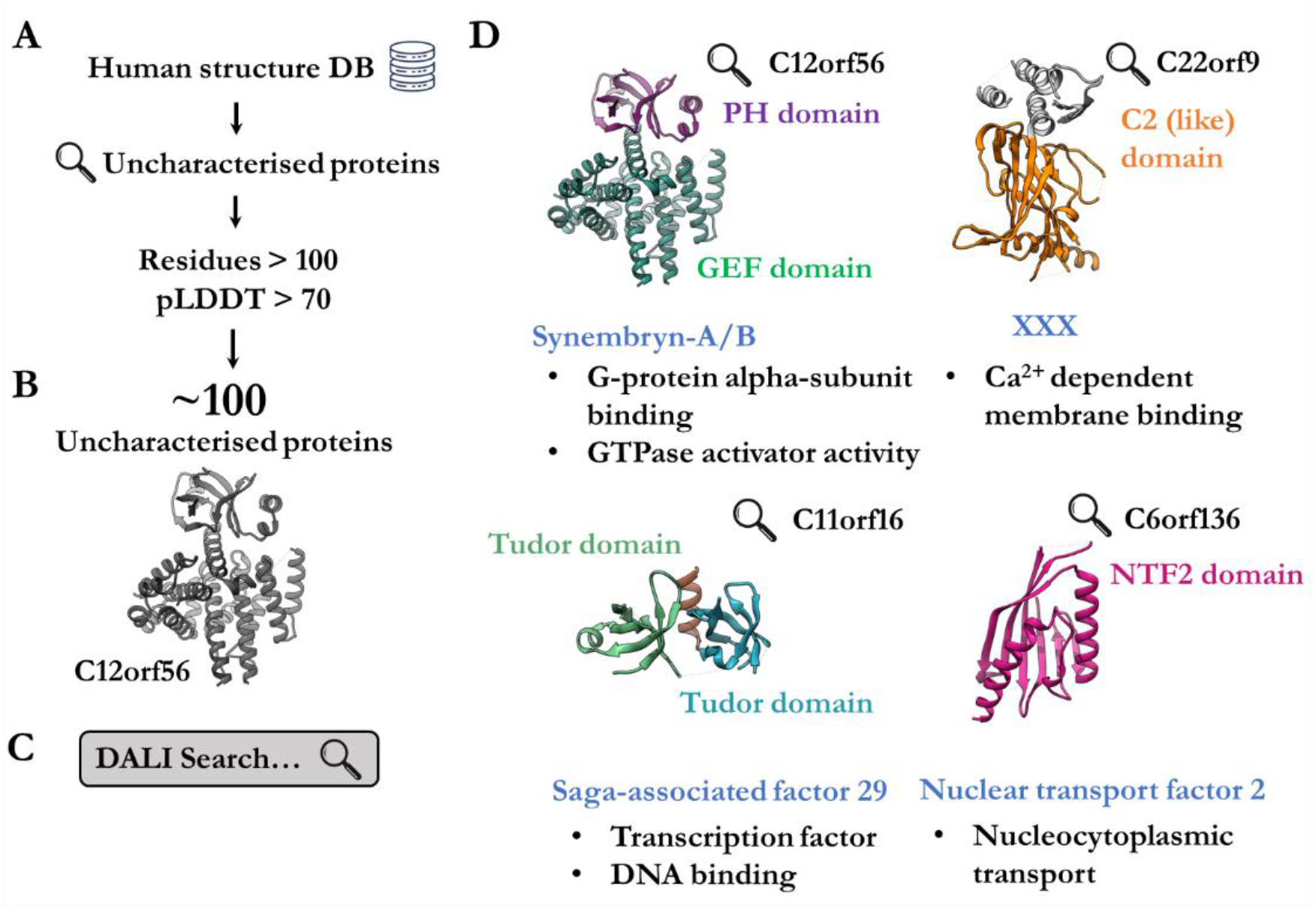
Identification of uncharacterised human proteins by fold-recognition. **a**. Uncharacterised human proteins were curated from the AlphaFold database. **b**. Low-confidence regions of AlphaFold models were excluded based on pLDDT criteria. **c**. All models were screened against the rest of the human AlphaFold database. **d**. Four examples (C12orf56, Q8IXR9; C22orf9, Q6ICG6; C11orf16, Q9NQ32; C6orf136, Q5SQH8) of uncharacterised proteins where fold-matching enabled the assignment of domain composition (labelled in various colours). Furthermore, homologs or similar proteins (blue label) provide insight into potential function (black dot points).

From these analyses, we highlight four notable examples of uncharacterised structures which met the criteria and yielded insight into potential function (**Fig 2d**). One of these, C12orf56, appears to be a previously unknown GTPase activator protein. When compared to its homolog Ric-8A^24^, the PH domain appears to sterically occlude binding of Gα proteins and may result in a potentially autoinhibited conformation. Previously, the identity of this protein was most likely obscured in sequence-based approaches due to the abnormally large loop insertion in the PH domain (**Fig 2d**). Similarly, a putative nuclear import factor (NFT) with strong homology to NFT2 was identified. These examples demonstrate the utility of structure-guided curation and annotation of uncharacterised proteins.

Lastly, in an attempt to employ structure matching to assign domain composition, we searched representative protein domains against the *Staphylococcus aureus* foldome (the smallest available foldome) to score unknown and known proteins according to their tertiary structure similarity (**Fig 3a**). These representative structures comprise the entire trRosetta Pfam library and curated exemplar structures from the PDB for Pfam entries not modelled by trRosetta. A remaining 19% of Pfam entries were excluded (fewer than 50 residues, absence of PDB entry). Here, we employed a GPU-acclerated fold recognition software, SA Tableau search^25^, to expedite the large comparison. The analysis outputs a ranked list of all proteins which match the query domain (**Supp Data 3**). As such these serve as first-pass approximation of structure-assigned domain composition for the *S. aureus* proteome. As an example, we selected entries of uncharacterised proteins within *S. aureus* and assigned these to Pfam classes based on the DALI results (**Fig 3b, c**). We note significant redundancy between Pfam groups with respect to structural homology.

**Figure 3.**
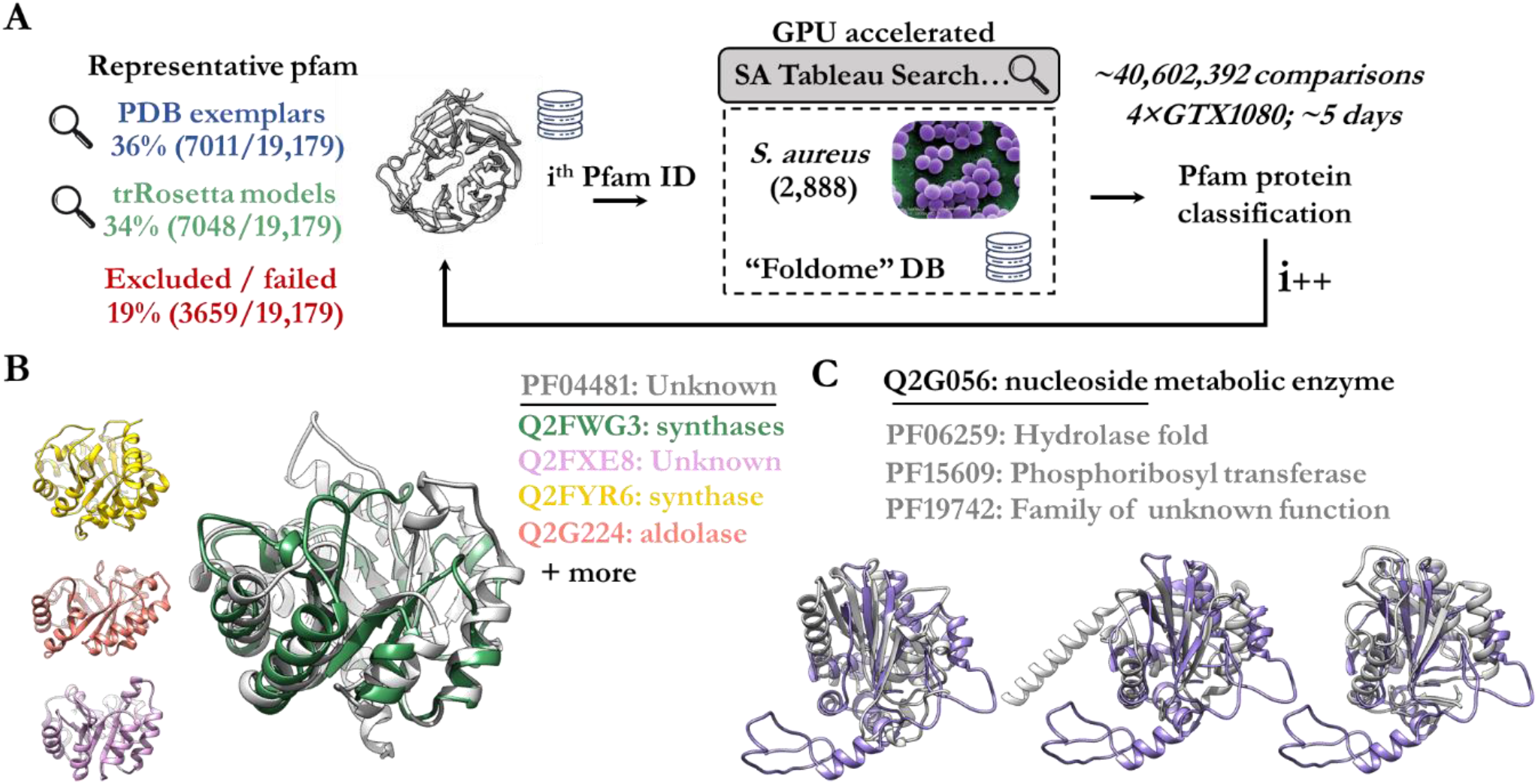
Proteome-wide search and classification of Pfam groups. **a**. Overview of analysis, firstly search models were curated from either trRosetta or the PDB that represent each Pfam classification. These were each searched against the entire *S. aureus* foldome to identify matches. The final output was filtered by expectation value. **b**. Select examples of *S. aureus* proteins that fell into Pfam category PF04481. This Pfam category previously had unknown role or function, now matches against proteins of known function. **c**. An uncategorised protein (Q2G056) is homologous to different Pfam groups (PF06259, PF15609, PF19742).

Exhaustive searches of predicted structures serve as a sensitive, but computationally expensive, domain-assignment tool for proteins which lack sequence annotations or where domain assignment has not been successful using sequence-based approaches. Owing to prohibitively time-consuming computational limitations, it was not feasible for us to search the entire foldome across all organisms. The dedication of high-performance computing resources to the remaining proteomes, or particularly subsets that are still unknown, may be merited. Similarly, the AlphaFold database would benefit from application of other structure-based classification methods (such as adaptations to classification schemes of SCOPe^26^ or ECOD^27^). The curated subset of PDB entries used for DALI searches are available as a resource to expedite efforts by others (**Supp Data 3**) and supplement the trRosetta Pfam models (publicly available: http://ftp.ebi.ac.uk/pub/databases/).

Finally, it is our perspective that adoption of the above analyses among structural biologists may be beneficial. Common practice of firstly searching for related folds before beginning a project, may accelerate investigations and improve the likelihood of success. For example, one might first generate an accurate structural prediction of the target molecule, then search this against larger foldome databases (perhaps via webserver) to gain insight into function and putative mechanism. In this way, before experimentation, previously obtained knowledge of function can provide rationale, guide inquiry and minimise unnecessary or resource intensive efforts – saving time and money. Likewise, the identification of homologs in model organisms may facilitate parallel studies in vivo or in situ.

Overall, the efficacy of structure mining depends on the accuracy of predicted models. Currently, this is dependent on MSAs for the detection of evolutionary covariance, however new single-sequence structure prediction methods are emerging that do not rely on sequence alignment^28^. Currently, the extent and quality of the MSA will affect the quality of AlphaFold/RoseTTAFold predictions and thus the quality of search results. Notably, protein families with extensive primary sequence conservation may not benefit from structure-guided mining, as existing techniques are likely sufficiently sensitive, as well as being far quicker and more computationally efficient. As such, protein folds that are structurally conserved but have poor overall sequence conservation may represent ideal targets for structure-based mining.

## Author contributions

Both authors conceived and developed the study. CBJ performed data analysis, bioinformatic investigation and produced all figures. Both authors co-wrote the draft, performed revisions and editing, interpreted findings, and reviewed the literature.

## Acknowledgements

CBJ acknowledges helpful discussion and feedback from Dr. Bradley A. Spicer and Dr. Andrew Ellisdon. JCW acknowledges funding from the Australian Research Council and the National Health and Medical Research Council.

## Materials and Methods

### Generation and acquisition of AlphaFold models

All AlphaFold models were obtained from the EMBL EBI database (https://alphafold.ebi.ac.uk/) for each available model organism. For any searches where existing models were not available from the PDB, these were generated using AlphaFold hosted through ColabFold^29^. A regex search of PDB metadata for ‘uncharacterised’ was used to curate a subset of uncharacterised or unknown proteins in the human foldome. Atomic coordinates for these files were subsequently discarded if their pLDDT score was less than 70. Remaining models which possessed fewer than 100 residues were discarded.

### Construction of local DALI search engine and database

All DALI searches were performed using DaliLite^3^ (v5; available from http://ekhidna2.biocenter.helsinki.fi/dali/) on one of two Linux workstations equipped with 16-core or 20-core Intel i7 CPU and 128 Gb of DDR4 RAM. DALI database was generated as described in the DALI manual. Briefly, for every AlphaFold model a randomised four character internal “PDB” code was generated and associated with the model (*unique_identifiers*.*txt*). Subsequently, all models were imported and converted to DALI format to enable structure all-versus-all searches. Individual proteomes were isolated as separate lists of entries or combined, to enable independent or grouped searches.

### Construction of local SA Tableau (GPU accelerate) search engine and database

All SA Tabelau searches were performed on the same Linux workstations as for the DALI searches; however jobs were split equally into four sets and each set was executed on a single Nvidia GTX1080 GPU. Since SA Tableau was originally written and compiled^25^ for outdated CUDA architectures, we re-compiled SA Tableau under modern CUDA (v8.0) gcc (v4.9.3) and g++ (v5.4.0). To run SA Tableau, it was necessary to create a conda environment with python2.7, where numpy (v1.8.1) and biopython (v1.49) were found to properly execute and run the original code. SA Tableau databases and distance matrices were calculated with ‘*buildtableauxdb*.*py*’ and combined into ASCII format with ‘*convdb2*.*py*’ (available from http://munk.cis.unimelb.edu.au/~stivalaa/satabsearch/ at the date of publication). SA Tableau results were sorted and selected based on expectation value with a cut-off of 1×10^−4^.

### Proteome-wide assignment of Pfam

In order to search the entire Pfam classification against a structural proteome database, we used the GPU-accelerated SA Tableau search algorithm to expedite the search process. Furthermore, we selected to search only the *S. aureus* foldome as this represents the smallest and therefore most computationally inexpensive example proteome. All Pfam classifications were represented by structure in one of two ways. Firstly, trRosetta models for ∼7,000 Pfam classifications were recently produced and used without modification. Secondly, of the remaining 60% of entries, we used the ProtCID database^30^ (http://dunbrack2.fccc.edu/ProtCiD/default.aspx) to link Pfam IDs to known protein structures in the PDB. These exemplar structures from the PDB were downloaded and single chains were extracted from each model (that is, only a single copy of each domain was considered). Domain boundaries and chain IDs defined by ProtCID were used to discard unrelated chains and residues that did not pertain to the particular Pfam classification in question. Finally, the trRosetta and exemplar structures were searched against the entire *S. aureus* foldome.

## Supplementary

**Supplementary Figure 1.**
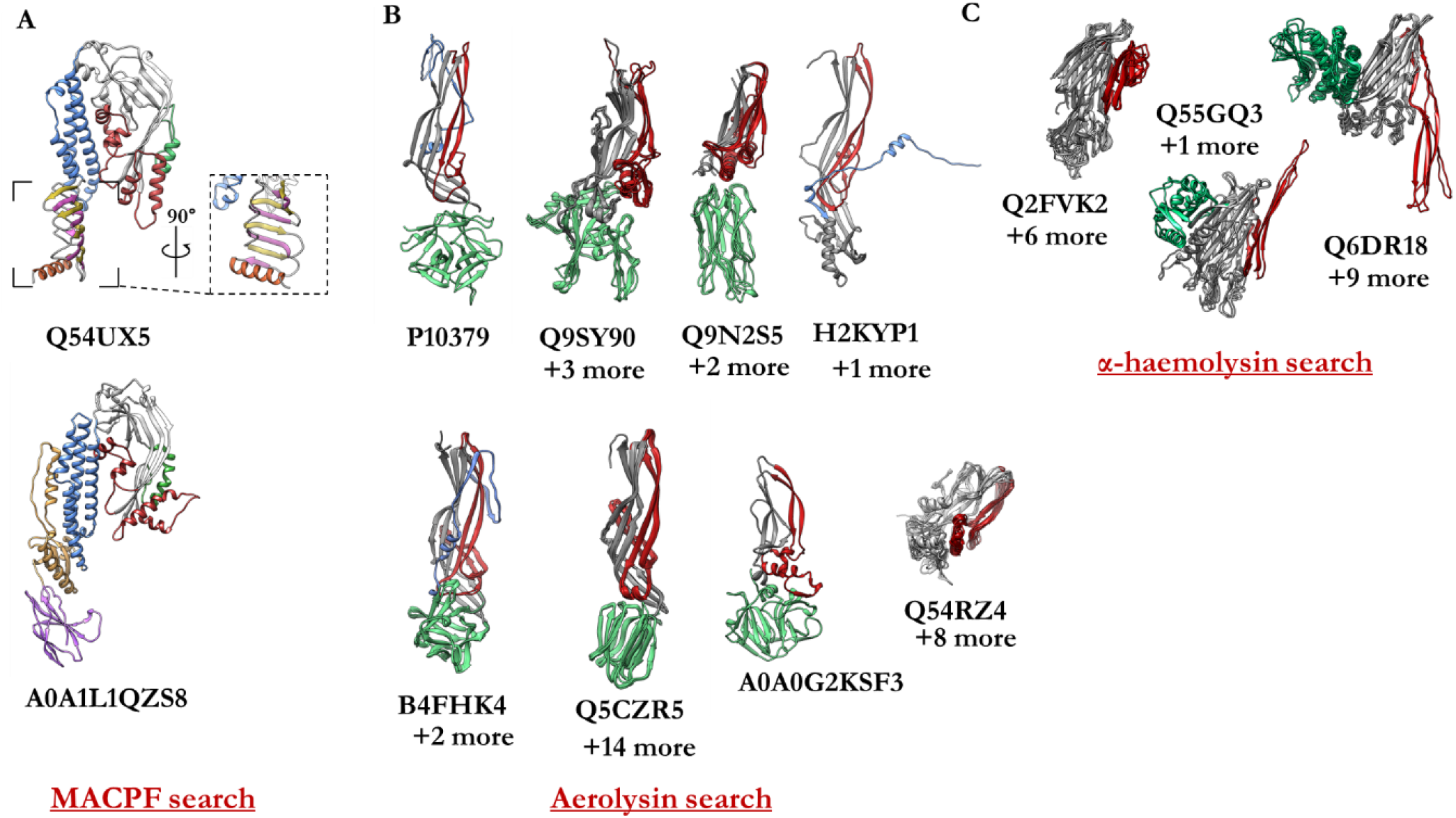
Select examples of newly identified PFPs. **a**. Identified MACPFs from slime mold and zebrafish. **b**. Various β-PFPs with aerolysin like pore forming domains which resemble aerolysin, lysenin, epsilon toxin, monalysin and LSL. Observed in numerous organisms including drosophila, *C. elegans*, zebrafish, yeast, among others. **c**. Several novel α-haemolysin-like proteins identified in *S. aureus* and plants.

**Supplementary Data 1, 2, 3** – see attached zip files.

**Supplementary Table 1** – see attached file – Curated lists of pore forming proteins identified by DALI search of AlphaFold database, organised by query.

## References

1. Andreeva, A. & Murzin, A. G. Structural classification of proteins and structural genomics: New insights into protein folding and evolution. Acta Crystallogr. Sect. F Struct. Biol. Cryst. Commun. 66, 1190–1197 (2010).

2. Andreeva, A., Kulesha, E., Gough, J. & Murzin, A. G. The SCOP database in 2020: Expanded classification of representative family and superfamily domains of known protein structures. Nucleic Acids Res. 48, D376–D382 (2020).

3. Holm, L. Dali server:conservation mapping in 3D. Nucleic Acids Res. 38, 545–549 (2010).

4. Rosado, C. J. et al. A common fold mediates vertebrate defense and bacterial attack. Science (80-.). 317, 1548–1551 (2007).

5. Jumper, J. et al. Highly accurate protein structure prediction with AlphaFold. Nature (2021). doi:10.1038/s41586-021-03819-2

6. Baek, M. et al. Accurate prediction of protein structures and interactions using a three-track neural network. Science (80-.). eabj8754 (2021). doi:10.1126/science.abj8754

7. Tunyasuvunakool, K. et al. Highly accurate protein structure prediction for the human proteome. Nature (2021). doi:10.1038/s41586-021-03828-1

8. Artificial intelligence in structural biology is here to stay. Nature 595, 625–626 (2021).

9. Porta-Pardo, E., Ruiz-Serra, V. & Valencia, A. The structural coverage of the human proteome before and after AlphaFold. bioRxiv 2021.08.03.454980 (2021). doi:10.1101/2021.08.03.454980

10. Uhlén, M. et al. Tissue-based map of the human proteome. Science (80-.). (2015). doi:10.1126/science.1260419

11. Behan, F. M. et al. Prioritization of cancer therapeutic targets using CRISPR–Cas9 screens. Nature (2019). doi:10.1038/s41586-019-1103-9

12. Reddy, A. et al. Genetic and Functional Drivers of Diffuse Large B Cell Lymphoma. Cell (2017). doi:10.1016/j.cell.2017.09.027

13. Morgens, D. W., Deans, R. M., Li, A. & Bassik, M. C. Systematic comparison of CRISPR/Cas9 and RNAi screens for essential genes. Nat. Biotechnol. (2016). doi:10.1038/nbt.3567

14. Oughtred, R. et al. The BioGRID database: A comprehensive biomedical resource of curated protein, genetic, and chemical interactions. Protein Sci. 30, 187–200 (2021).

15. Ruan, J., Xia, S., Liu, X., Lieberman, J. & Wu, H. Cryo-EM structure of the gasdermin A3 membrane pore. Nature 557, 62–67 (2018).

16. Ding, J. et al. Pore-forming activity and structural autoinhibition of the gasdermin family. Nature 535, 111–116 (2016).

17. Ni, T., Harlos, K. & Gilbert, R. Structure of astrotactin-2: A conserved vertebrate-specific and perforin-like membrane protein involved in neuronal development. Open Biol. 6, (2016).

18. Van der Verren, S. E. et al. A dual-constriction biological nanopore resolves homonucleotide sequences with high fidelity. Nat. Biotechnol. 38, 1415–1420 (2020).

19. Goyal, P. et al. Structural and mechanistic insights into the bacterial amyloid secretion channel CsgG. Nature 516, 250–3 (2014).

20. Brinkerhoff, H., Kang, A. S. W., Liu, J., Aksimentiev, A. & Dekker, C. Infinite re-reading of single proteins at single-amino-acid resolution using nanopore sequencing. bioRxiv (2021).

21. Howorka, S. & Siwy, Z. S. Reading amino acids in a nanopore. Nat. Biotechnol. 38, 159–160 (2020).

22. Nivala, J., Mulroney, L., Li, G., Schreiber, J. & Akeson, M. Discrimination among protein variants using an unfoldase-coupled nanopore. ACS Nano 8, 12365–12375 (2014).

23. Ouldali, H. et al. Electrical recognition of the twenty proteinogenic amino acids using an aerolysin nanopore. Nature Biotechnology 38, 176–181 (2020).

24. McClelland, L. J. et al. Structure of the G protein chaperone and guanine nucleotide exchange factor Ric-8A bound to Gαi1. Nat. Commun. 11, (2020).

25. Stivala, A. D., Stuckey, P. J. & Wirth, A. I. Fast and accurate protein substructure searching with simulated annealing and GPUs. BMC Bioinformatics 11, 446 (2010).

26. Chandonia, J. M., Fox, N. K. & Brenner, S. E. SCOPe: Classification of large macromolecular structures in the structural classification of proteins - Extended database. Nucleic Acids Res. 47, D475–D481 (2019).

27. Cheng, H. et al. ECOD: An Evolutionary Classification of Protein Domains. PLoS Comput. Biol. 10, (2014).

28. Chowdhury, R. et al. Single-sequence protein structure prediction using language models from deep learning. bioRxiv 2021.08.02.454840 (2021). doi:10.1101/2021.08.02.454840

29. Mirdita, M., Ovchinnikov, S. & Steinegger, M. ColabFold - Making protein folding accessible to all. bioRxiv 2021.08.15.456425 (2021). doi:10.1101/2021.08.15.456425

30. Xu, Q. & Dunbrack, R. L. ProtCID: a data resource for structural information on protein interactions. Nat. Commun. 11, 711 (2020).

