## Supplementary Data 1 for "Mining folded proteomes in the era of accurate structure prediction": README.txt

######

Charles Bayly-Jones & James C. Whisstock 
"Mining folded proteomes in the era of accurate structure prediction" 2021
bioRxiv: 




######

--------------------
Supplementary Data 1
--------------------

#Included:
README.txt - this file
unique_identifiers.txt - lookup table mapping internal unique identifiers (these DO NOT correspond to RCSB PDB codes) to corresponding alphafold models
	e.g. 'grep k0we unique_identifiers.txt' => foldomeDB/human/AF-P30085-F1-model_v1.pdb.gz
	Either download alphafold proteome from https://alphafold.ebi.ac.uk/download OR access individual model from Uniprot (all alphafold models contain 
	uniprot ID directly after 'AF-')
raw_search_results/ - directory: raw results file of DALI search including all matches for search against given target.
