## Supplementary Data 2 for "Mining folded proteomes in the era of accurate structure prediction": README.txt

######

Charles Bayly-Jones & James C. Whisstock 
"Mining folded proteomes in the era of accurate structure prediction" 2021
bioRxiv: 




######

--------------------
Supplementary Data 2
--------------------

#Included:
- README.txt - this file
- unique_identifiers.txt - lookup table mapping internal unique identifiers (these DO NOT correspond to RCSB PDB codes) to corresponding alphafold models
	e.g. 'grep k0we unique_identifiers.txt' => foldomeDB/human/AF-P30085-F1-model_v1.pdb.gz 
	Either download alphafold proteome from https://alphafold.ebi.ac.uk/download OR access individual model from Uniprot (all alphafold models contain uniprot ID directly after 'AF-')
- search_results/ - directory containing all raw result files from DALI search. Each file is named according to the AlphaFold model (by Uniprot ID) used as the probe (i.e. search template). Each file contains typical DALI results, with internal unique identifiers (use look up table).
