## Supplementary Data 3 for "Mining folded proteomes in the era of accurate structure prediction": README.txt

######

Charles Bayly-Jones & James C. Whisstock 
"Mining folded proteomes in the era of accurate structure prediction" 2021
bioRxiv: 




######

--------------------
Supplementary Data 3
--------------------

README.txt - this file.
SATableau_search/ - raw output files for each search. File is named according to Pfam ID and trRosetta model used for search. Results are Uniprot IDs.
filtered.sort.output - Sorted text file containing raw results of search filtered to have an expectation value larger than 1e-3
filtered_output.xlsx - Same as above, in excel format.
PF04481.py - UCSF chimera (v1.13) session file containing superposition of Pfam PF04481 matches against AlphaFold models (uniprot ID).
Q2G056.py - UCSF chimera (v1.13) session file containing superposition of uniprot Q2G056 matches to trRosetta Pfam models.

trRosetta models available at: 
http://ftp.ebi.ac.uk/pub/databases/Pfam/releases/Pfam34.0/structure_models/
